# Chromatin accessibility dynamics reveal novel functional enhancers in *C. elegans*

**DOI:** 10.1101/088732

**Authors:** Aaron C. Daugherty, Robin Yeo, Jason D. Buenrostro, William J. Greenleaf, Anshul Kundaje, Anne Brunet

## Abstract

Chromatin accessibility, a crucial component of genome regulation, has primarily been studied in homogeneous and simple systems, such as isolated cell populations or early-development models. Whether chromatin accessibility can be assessed in complex, dynamic systems *in vivo* with high sensitivity remains largely unexplored. In this study, we use ATAC-seq to identify chromatin accessibility changes in a whole animal, the model organism *C. elegans*, from embryogenesis to adulthood. Chromatin accessibility changes between developmental stages are highly reproducible, recapitulate histone modification changes, and reveal key regulatory aspects of the epigenomic landscape throughout organismal development. We find that over 5,000 distal non-coding regions exhibit dynamic changes in chromatin accessibility between developmental stages, and could thereby represent putative enhancers. When tested *in vivo*, several of these putative enhancers indeed drive novel cell-type-and temporal-specific patterns of expression. Finally, by integrating transcription factor binding motifs in a machine learning framework, we identify EOR-1 as a unique transcription factor that may regulate chromatin dynamics during development. Our study provides a unique resource for *C. elegans*, a system in which the prevalence and importance of enhancers remains poorly characterized, and demonstrates the power of using whole organism chromatin accessibility to identify novel regulatory regions in complex systems.

## Introduction

Chromatin accessibility represents an essential level of genome regulation and plays a pivotal role in many biological and pathological processes, including development, tissue regeneration, aging, and cancer (Simon et al. 2014; Stergachis et al. 2013; Tsompana et al. 2014). However, most genome-wide chromatin accessibility studies to date have been in relatively simple systems, including cultured or purified cells, and early embryos (Lara-Astiaso et al. 2014; Thomas et al. 2011; Wang et al. 2012; West et al. 2014). Assessing chromatin accessibility directly in complex systems composed of multiple cell types could allow for high-throughput discovery of regulatory regions whose activity are restricted to rare or undefined sub-populations of cells. This is particularly relevant for enhancers, which are thought to be highly cell-type-and temporally-specific (Ren et al. 2015).

The primary limitation for studying chromatin accessibility in complex systems is that most assays lack the sensitivity and precision to detect regions active only in rare sub-populations of cells, or require so many cells that precise temporal synchronization of samples is impractical. However, the Assay for Transposase Accessible Chromatin using sequencing (ATAC-seq) has been shown to assess native chromatin accessibility with high sensitivity and base pair resolution while requiring orders of magnitude less starting material than other assays (Buenrostro et al. 2013). This approach has been used successfully in cultured or purified cells even down to single cells, though such low input relies heavily on existing knowledge (Buenrostro et al. 2015; Cusanovich et al. 2015). Rather than purifying specific cell types, we wondered whether ATAC-seq could be sensitive enough to detect subtle changes in chromatin accessibility in complex mixtures of tissue, and, in so doing, uncover novel biological insights that would have otherwise been obscured.

The nematode *Caenorhabditis elegans* is a particularly powerful model to study chromatin accessibility in a complex system and potentially identify novel regulatory regions. *C. elegans* has highly synchronous life stages, as well as consistent and well characterized cellular composition throughout each stage of development (Sulston et al. 1983). Rapid transgenesis and transparency (Mello et al. 1995) also make *C. elegans* an ideal system to efficiently validate genomic regions of functional importance and visualize tissue- or cell-specificity (Harfe et al. 1998; Jantsch-Plunger et al. 1994; Lei et al. 2009). While there exist some preliminary reports on chromatin states in *C. elegans* (Evans et al. 2016; Gerstein et al. 2010; Hsu et al. 2015; Liu et al. 2011b; Shi et al. 2009; Valouev et al. 2008), high-resolution, genome-wide chromatin accessibility maps throughout development have not yet been reported. In this study, we show that studying high-resolution chromatin accessibility dynamics in synchronized *C. elegans* populations allows us to characterize highly reproducible changes in chromatin accessibility between developmental stages and to identify functional temporal- and tissue-specific novel enhancers *in vivo*. Our study provides a unique resource for defining *C. elegans* regulatory regions as well as a guide for the interpretation of chromatin structure in complex multi-tissue systems *in vivo*.

## Results

### High-resolution chromatin accessibility profiles from three *C. elegans* life stages

To sensitively measure high-resolution chromatin accessibility at different life stages in *C. elegans*, we used the Assay for Transposase Accessible Chromatin using sequencing (ATAC-seq). We optimized the ATAC-seq protocol for *C. elegans* by including a step of native nuclei isolation by mechanical homogenization before the transposition step (see Methods and Supplemental Extended Protocol). The low input requirements of ATAC-seq (several orders of magnitude less than standard ChIP-seq (Furey 2012)) allowed us to grow *C. elegans* in standard conditions (i.e. plates, whereas most high-throughput assays in *C. elegans* require growth in liquid) and to generate three independent biological replicates that are tightly synchronized at three key life stages: early embryo, larval stage 3 (L3), and young adults, thereby limiting variation within stages (see Methods) (Fig. 1A). We generated and sequenced ATAC-seq libraries, as well as an input control, to a median depth of over 17 million unique, high-quality mapping reads per sample (Supplemental Table 1). The insert size distribution of each *C. elegans* ATAC-seq experimental library displays the stereotypical 147 bp periodicity (Supplemental Fig. 1A), consistent with the expected nucleosome occupancy of chromatin. We designed a computational framework to integrate the input control (Supplemental Fig. 1B) and emphasize single base pair resolution (Fig. 1B), resulting in the identification of 13,000 to 27,000 high-confidence, accessible peaks per developmental stage and more than 30,000 consensus ATAC-seq peaks found in at least one of the three stages (Supplemental Tables 2 and 3, see Methods). The high correlation of ATAC-seq signal between each of the 3 biological replicates (Spearman’s rho > 0.837) demonstrates the high reproducibility of this approach (Fig. 1C). The ability to cluster samples by their developmental stage also shows that chromatin accessibility is strikingly different

**Table 1.** Transgenic reporter lines generated for validation of ATAC-seq peaks as functional enhancers. Summary of the number of independent transgenic strains generated to assess functional activity of putative enhancer regions identified by ATAC-seq dynamics. The number of independent strains that exhibited consistent spatiotemporal expression patterns of GFP with inserted ATAC-seq peaks in the native genomic orientation, reverse orientation, and with a nearby a flanking region, is reported here.

| Table 1. | 5' → 3' Orientation | 3' → 5' Orientation | Negative Control (flanking) |
| --- | --- | --- | --- |
| Putative Gene | # GFP+ / total lines | # GFP+ / total lines | # GFP+ / total lines |
| C54G6.3 | 8/8 | 2/2 | 0/3 |
| *nhr-25* | 8/8 | 3/3 | 0/8 |
| *swip-10* | 8/8 | 2/2 | 0/1 |
| *mlt-8* | 11/11 | 5/5 | 0/3 |
| *gei-13* | 10/11 | 6/6 | 0/1 |
| No-insert control | 0/3 | - | - |

**Figure 1.**
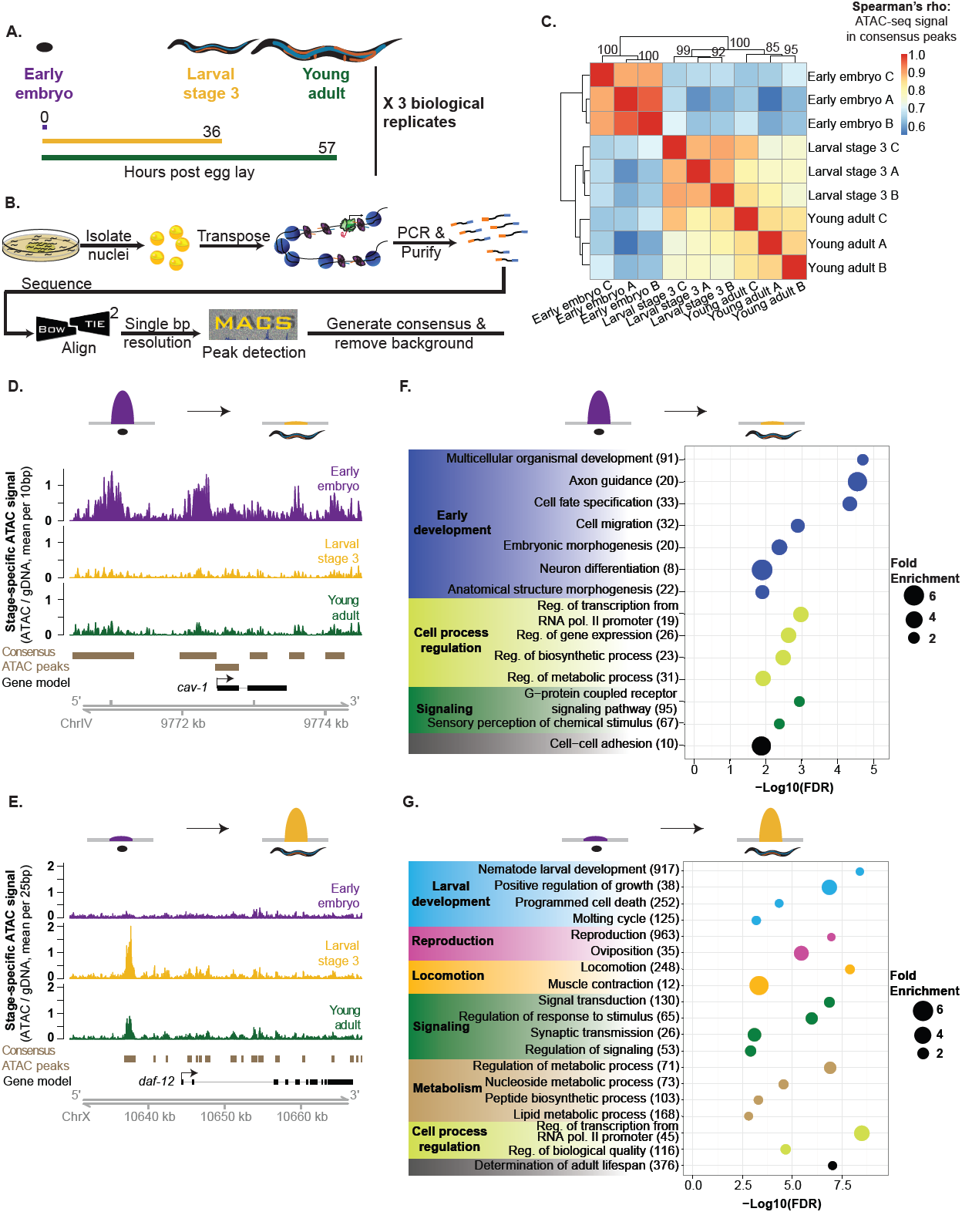
ATAC-seq in whole *C. elegans* captures chromatin accessibility dynamics across three life-stages. (*A*) Three independent biological replicates each consisting of tightly temporally synchronized *C. elegans* were used for ATAC-seq. Hours post egg-lay are at 20°C. (*B*) *C. elegans* were flash frozen and nuclei were isolated before assaying accessible chromatin using transposons loaded with next-generation sequencing adaptors, allowing paired-end sequencing. A custom analysis pipeline emphasizing high-resolution signal and consistent peaks, as well as accommodating input control was developed to generate stage-specific and consensus (i.e. across stages) ATAC-seq peaks. (*C*) ATAC-seq signal within consensus ATAC-seq peaks was compared between all samples using Spearman’s rho to cluster samples. Replicate batches are noted as letters following the stage. (*D,E*) Comparison of ATAC-seq signal (normalized by total sequencing depth) between all three stages at a region that decreases (D) or increases (E) in accessibility during development. (*F,G*) Genes that lose accessibility between embryo and larval stage 3 (L3) are enriched for early development functions (F), while genes that gain accessibility are enriched for larval-related functions (G); all calculations and genes lists are from GOrilla and the number of genes enriched in each term are listed in parentheses.between these three life stages (Fig. 1C). These differences between developmental stages are likely due to both changes in accessibility within cells, as well as the organisms’ changing cellular composition throughout development. Together, these results indicate that reproducible high-resolution chromatin accessibility can be obtained from low amounts (at least an order of magnitude less than standard histone ChIP-seq or DNase-seq) of complex, multi-tissue samples.

To investigate the changes in chromatin accessibility between life stages, we identified and characterized the ATAC-seq peaks that significantly changed accessibility between early embryo and L3 (12,193 peaks, Supplemental Fig. 1C) and between L3 and young adult (783 peaks, Supplemental Fig. 1D) (FDR < 0.05) (see Methods). The larger number of differentially accessible peaks (both decreased and increased) observed during the transition from early embryo to L3 versus L3 to young adult could not simply be explained by differences in sequencing depth (see Methods) and is likely due to the massive changes in cell number and tissue composition that occur during this transition (Byerly et al. 1976).

An example of a decrease in chromatin accessibility from embryo to L3 can be seen in the promoter region of the *cav-1* gene, which is expressed during embryogenesis but not larval development (Parker et al. 2009) (Fig. 1D). Conversely, several ATAC-seq peaks drastically increase from embryo to L3 in the promoter and regions upstream of the *daf-12* gene, which is a key regulator of stage-specific developmental programs particularly at L3 (Antebi et al. 1998; Antebi et al. 2000) (Fig. 1E). Confirming these specific examples, the most enriched gene ontology (GO) terms for genes with decreased chromatin accessibility from embryo to L3 include early-development terms such as embryonic morphogenesis and cell fate specification (Fig. 1F and Supplemental Table 4), while the most enriched terms for genes with increased chromatin accessibility from embryo to L3 include larval development and locomotion (Fig. 1G and Supplemental Table 5). Similarly, we observe strong enrichments of GO terms reflecting the major phenotypic changes occurring between L3 and adult, including terms like larval development and reproduction (Supplemental Fig. 1E, 1F and Supplemental Table 6 and 7).

Together, these results indicate that ATAC-seq in whole organisms can identify changes in DNA accessibility that represent key biological differences between stages, regardless of whether these changes are due to activation/repression of specific regions within a cell-type or to changes in cell-type composition.

### ATAC-seq as a single assay describes the epigenome

Accessible chromatin encompasses several key features of the epigenome, including active and poised regulatory regions. To verify that our ATAC-seq data correctly identifies regulatory regions throughout the epigenome, we used multiple histone modification ChIP-seq datasets from modENCODE (Ho et al. 2014; Li et al. 2009) and ChromHMM (Ernst et al. 2012) to build predictive models of the epigenome. ChromHMM is a hidden Markov model that classifies regions of the genome into chromatin states (e.g. heterochromatin) using the co-occurrence of multiple histone modifications from each life stage – in our case, ChIP-seq datasets characterizing 8 distinct histone modifications across all three stages (Supplemental Fig. 3A-E; Supplemental Tables 8-11) (see Methods). ATAC-seq peaks from all three stages were significantly enriched in active and poised regulatory chromatin states (e.g. promoter), as defined by this ChromHMM model, and significantly depleted in heterochromatic states (Fig. 2A; Supplemental Fig. 3B, 3C). ATAC-seq peaks were also enriched in H3K27me3-repressed regions, which is likely due to H3K27me3 marking inactive, yet accessible poised sites (Lei et al. 2015; Lorzadeh et al. 2016; Rada-Iglesias et al. 2011; Sen et al. 2016; Zentner et al. 2011; Zhang et al. 2012). ATAC-seq signal was correlated with individual active histone modifications at transcription start sites (Fig. 2B) and genome-wide (Supplemental Fig. 3F). Thus, ATAC-seq correctly identifies poised and active regulatory regions at both specific loci and genome-wide.

**Figure 2.**
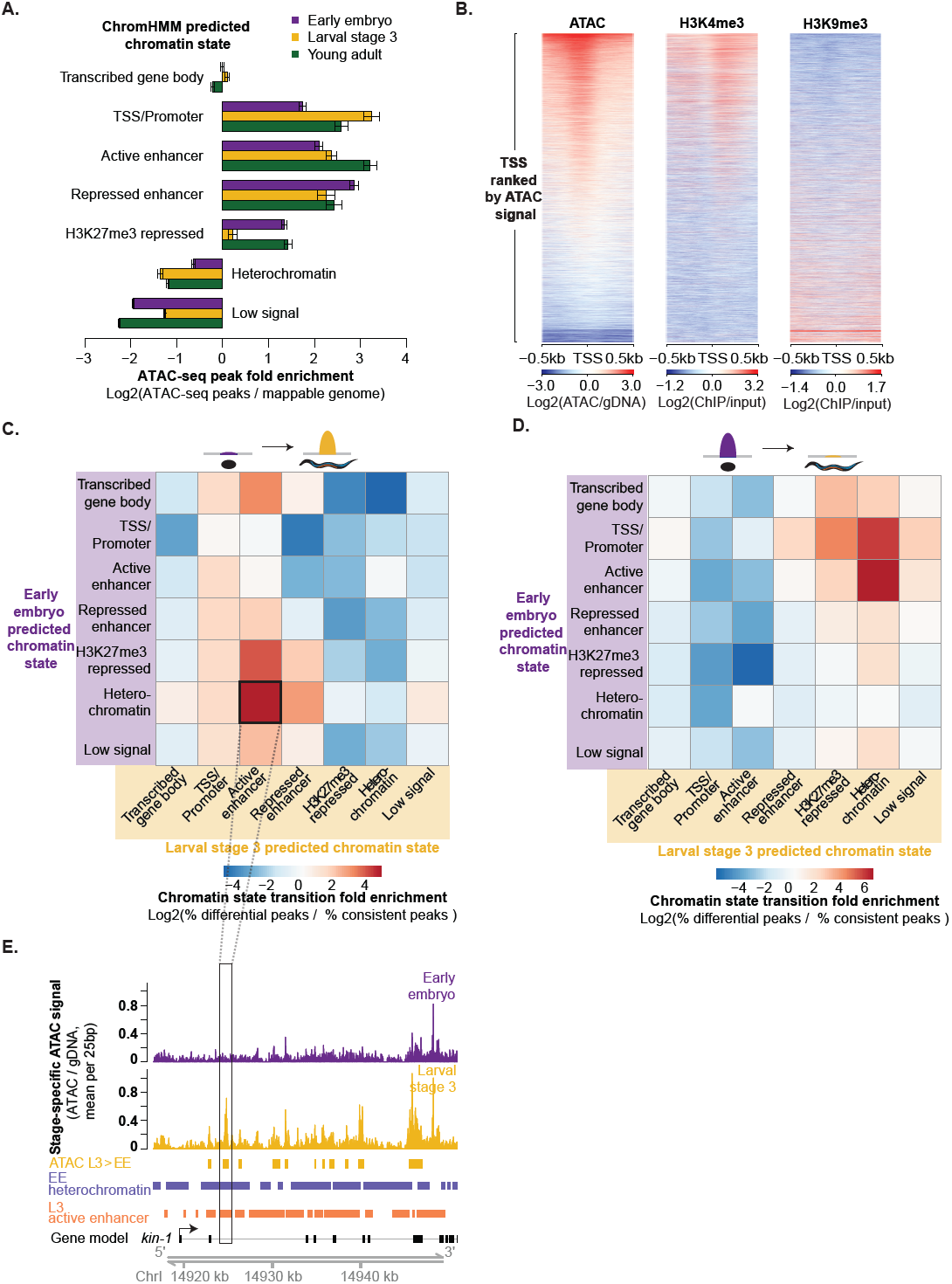
ATAC-seq as a single assay describes the epigenome. (*A*) Enrichment of stage specific ATAC-seq peaks in ChromHMM-predicted chromatin states relative to values expected by chance; significance derived via 10,000 bootstrapping iterations and error bars reflect 95% confidence intervals (p ≤ 1e-4, except for early embryo transcribed gene body (p = 0.633)). The decrease in enrichment of ATAC-seq peaks in H3K27me3-repressed regions in L3 compared to young adult and early embryo is likely due to the low sequencing depth of the L3 H3K27me3 ChIP-seq. (*B*) Larval stage 3 (L3) ATAC-seq signal at 19,899 previously defined transcription start sites (TSS +/- 0.5kb) is correlated with active histone modifications (H3K4me3) and anti-correlated with heterochromatin (H3K9me3) around the TSS. (*C,D*) Regions that increase in accessibility in ATAC-seq peaks are enriched for transitions from inactive chromatin states to active regulatory states (C), while regions that decrease in accessibility are enriched for transitions from active regulatory states to inactive chromatin states (D). (*E*) An example of an increase in chromatin accessibility overlapping with a transition from heterochromatin to a predicted active enhancer chromatin state. Multiple TSS have been noted for *kin-1*, but only the 5’-most is shown here for ease.

**Figure 3.**
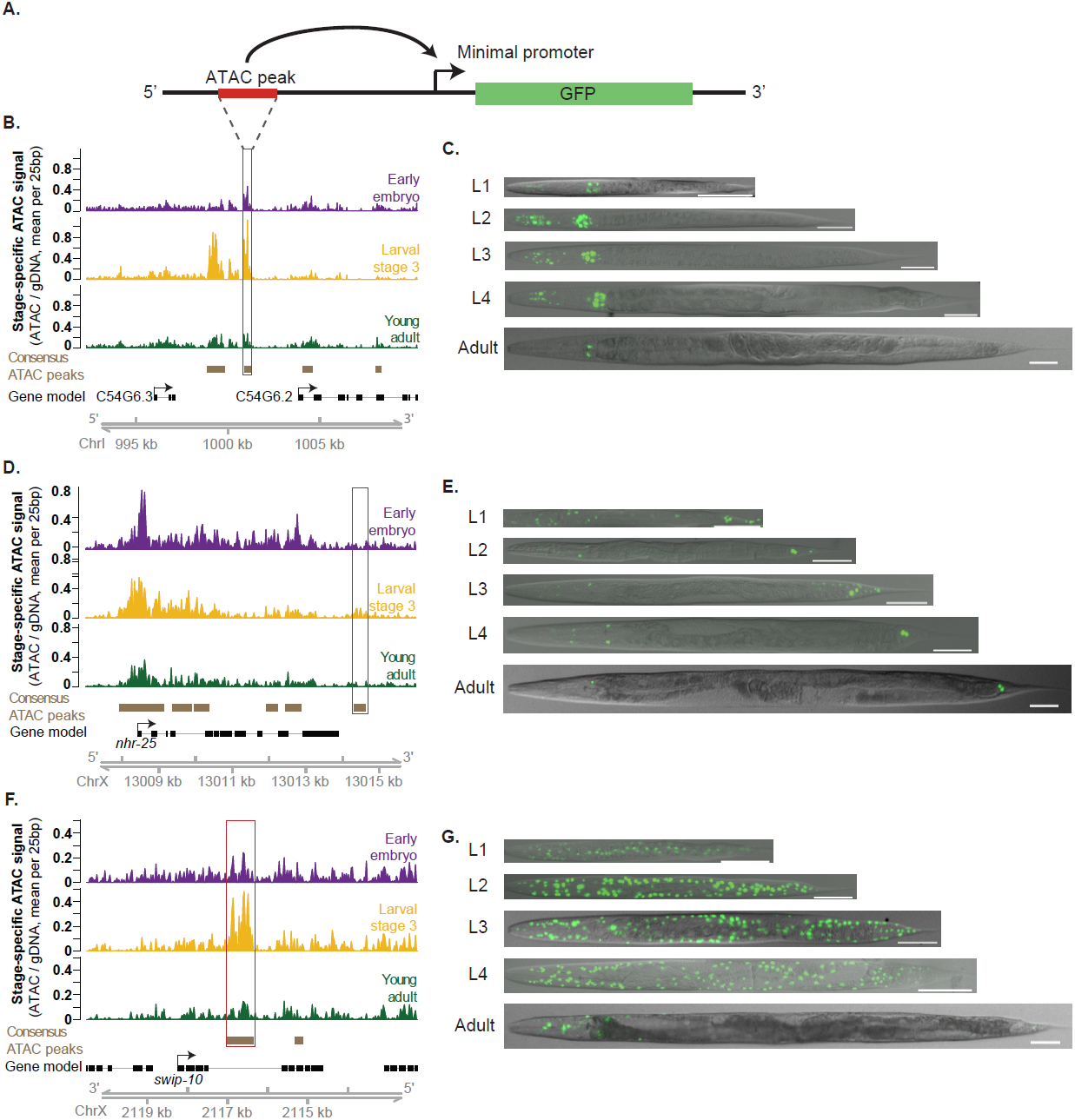
Dynamic ATAC-seq peaks identify functional enhancers with unique spatiotemporal specificity during ***C. elegans* development.** (A) Functional enhancer constructs used to generate *C. elegans* transgenic lines. Putative regulatory regions, or corresponding flanking regions for negative controls, are inserted upstream of a minimal promoter (*pes-10*) driving green fluorescent protein (GFP) localized to the nucleolus. (*B,D,F*) Distal (> 1 kb from a TSS) non-coding ATAC-seq peaks with the largest fold change in ATAC-seq signal between any two stages were screened for potential enhancer activity. The approximate regions tested near *C54G6.2* (*B*), *nhr-25* (*D*), and *swip-10* (*F*) are boxed in red. Note that for *swip-10* (F), the plot orientation is reversed for consistency with other plots. (*C, E*, G) Specific patterns of spatiotemporal enhancer activity in transgenic lines. Representative images of GFP expression in staged *C. elegans* transgenic lines are presented with a 50 μm scale bar. All images were straightened with ImageJ and are grayscale images with florescence overlaid.

To further assess the relevance of whole organism chromatin accessibility, we compared our ATAC-seq data to publicly available gene expression data (GRO-seq, and RNA-seq) (Gerstein et al. 2010; Gerstein et al. 2014; Hillier et al. 2009; Kruesi et al. 2013). Both GRO-seq (Supplemental Fig. 2A, 2B) and RNA-seq (Supplemental Fig. 2C, 2D) were positively correlated with ATAC-seq signal near the transcription start site (TSS). These observations support the relationship between chromatin accessibility at the TSS and gene expression, despite the numerous other layers of gene regulation (e.g. RNA stability).

Beyond simply identifying regulatory regions important for individual life stages, chromatin accessibility dynamics should highlight regulatory regions critical for transitions from embryo to larval stages, and from larval stages to adulthood. We examined whether genomic regions that showed accessibility changes from one life stage to another were enriched for specific chromatin state transitions. Regions that lost chromatin accessibility from embryo to L3 or from L3 to young adult were enriched for transitions from active regulatory chromatin states (especially predicted enhancers) to repressed or heterochromatic states (Fig. 2D; Supplemental Fig. 3G). Conversely, the regions that gained accessibility during development were enriched for transitions from inactive chromatin states to active regulatory states (again, especially predicted enhancers) (Fig. 2C, 2E; Supplemental Fig. 3H). Collectively these results show that ATAC-seq performed on an entire organism, even a complex multi-tissue adult, is sensitive enough to detect important global changes in chromatin structure that are consistent with predicted chromatin state transitions. Thus, ATAC-seq as a single assay constitutes an attractive alternative to performing multiple histone modification ChIP-seq experiments for identifying key regulatory regions, especially considering that ATAC-seq requires orders of magnitude less input than a single ChIP-seq.

### ATAC-seq identifies new distal regulatory regions that serve as tissue- and stage-specific enhancers in *C. elegans*

Enhancers are key regulators of temporal- and tissue-specific gene expression that play important and conserved functions during development (Ren et al. 2015; Sun et al. 2015). However, the identification and characterization of novel enhancers remains a challenge because they can be specifically active in rare cell populations, some of which may not have even been characterized (Heinz et al. 2015). The extent and functional importance of distal regulatory regions in the *C. elegans* genome has been particularly underexplored (Reinke et al. 2013). While several methods to identify potential enhancers genome-wide have recently been developed, these methods suffer from notable drawbacks (Arnold et al. 2013; Giresi et al. 2007; He et al. 2010; Mito et al. 2007; Visel et al. 2009; Wang et al. 2008; Zhu et al. 2015; Zhu et al. 2013). For example, the co-occurrence of multiple histone modifications (e.g. H3K27ac, H3K4me1) or RNA polymerase II (PolII) through ChIP-seq experiments lacks the sensitivity to detect enhancers active only in rare sub-populations of cells and the resolution to precisely identify the active enhancer region (Furey 2012). We therefore investigated whether whole-organism ATAC-seq could overcome these challenges and facilitate the identification of tissue- and stage-specific enhancers. Active and repressed enhancers, identified using the ChromHMM model and distinguished by the presence of H3K27ac (active enhancers) or H3K27me3 (repressed enhancers) (Supplemental Fig. 3A), were highly and significantly enriched in distal non-coding ATAC-seq peaks in all three stages (Supplemental Fig 4A). Furthermore, these distal non-coding ATAC-seq peaks were significantly more conserved than expected by chance (Supplemental Fig. 4B, 4C), a defining feature of enhancers (Pennacchio et al. 2006). ATAC-seq peaks also exhibited significant overlap with PolII transcription initiation complex binding at distal regions (p < 1e-323, one-sided Fisher’s exact test), a feature used previously to predict enhancers at a single stage (embryos) (Chen et al. 2013). These analyses support the notion that distal non-coding ATAC-seq peaks include enhancers.

**Figure 4.**
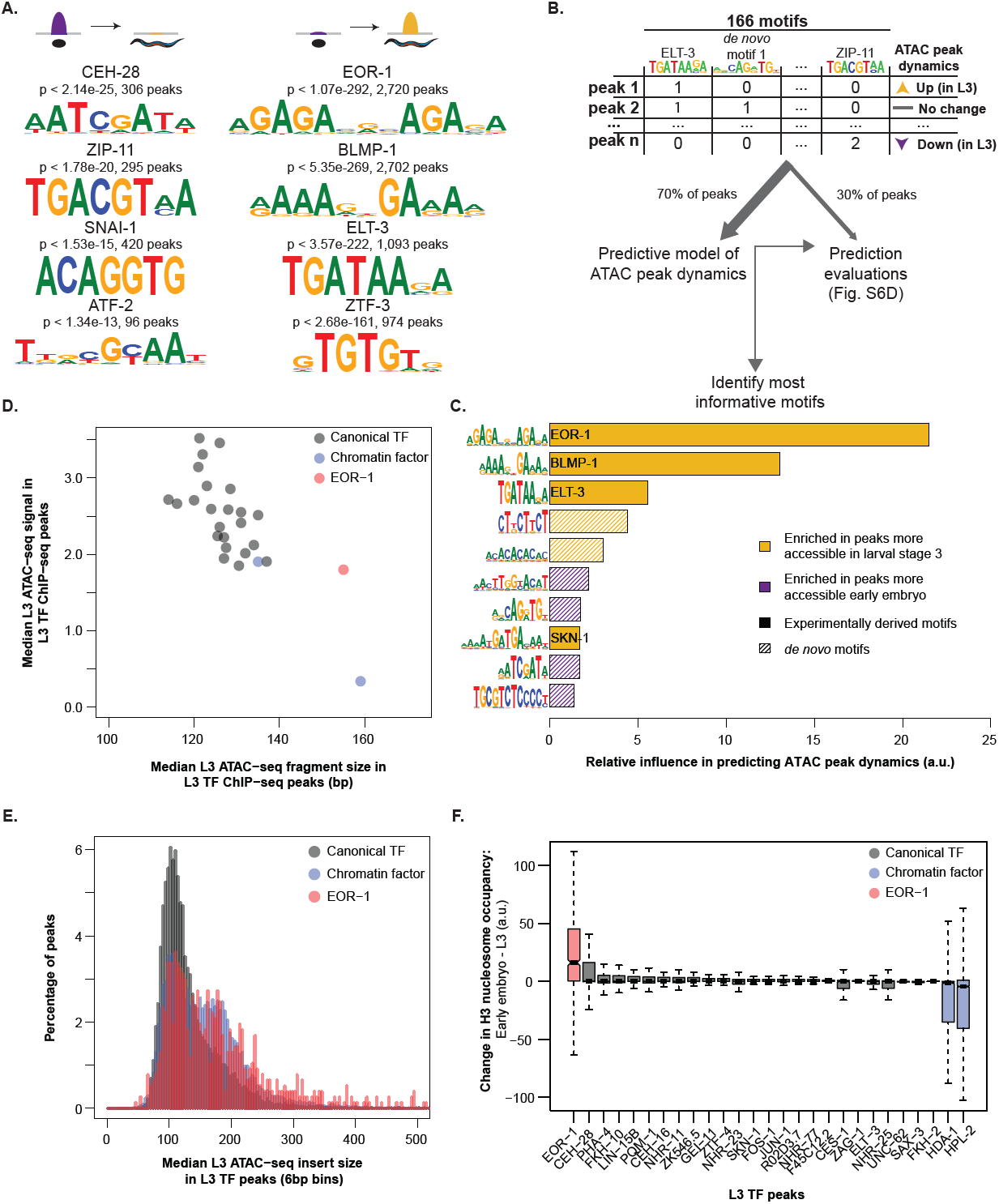
Motifs associated with increases in chromatin accessibility during development reveal key transcription factors with unique binding loci. (*A*) ATAC-seq peaks which decreased (left) or increased (right) accessibility between early embryo and L3 are enriched for previously identified transcription factor binding motifs; p-values are Benjamini-Hochberg corrected for multiple hypothesis testing. (*B*) The number of instances of previously identified as well as *de novo* motifs (see Fig. S6C) in each consensus ATAC-seq peak were used as features in a machine learning model to predict how each ATAC-seq peak changed between early embryo and L3 (increasing, decreasing, or no change). A training set (70% of all ATAC-seq peaks) was used to build the model, while the remaining held-out testing set was used to assess model quality (see Fig. S6D). (*C*) The relative influence of every motif from the machine learning model in Fig. 4B was quantified. Solid bars are previously defined motifs, while hashed bars are *de novo* identified motifs in dynamic ATAC-seq peaks. (*D*) The median L3 ATAC-seq signal and fragment length at the midpoint (+/- 50 bp) of L3 ChIP-seq peaks; box plots of the same data are in Fig. S7C, S7D. (*E*) Histograms of L3 ATAC-seq fragment size at the midpoint (+/- 50 bp) of L3 ChIP-seq peaks were calculated and normalized to percentages. Canonical TFs and chromatin factors were then aggregated and plotted. (*F*) The change in H3 nucleosome occupancy between early embryo and larval stage 3 at the midpoint of each L3 transcription factor ChIP-seq peak was calculated using Danpos and publicly available H3 ChIP-seq.

In *C. elegans*, a small number of functional enhancers have been experimentally mapped, including four enhancers in the upstream regulatory region of *hlh-1*, the *C. elegans* MyoD ortholog that regulates muscle development (Lei et al. 2009). ATAC-seq peaks overlap three of these four regions, and despite its lack of statistical significance, the fourth region still exhibits noticeable ATAC-seq signal in embryos (Supplemental Fig. 4D). Given that *hlh-1* is exclusively expressed in muscle (Krause et al. 1994), a tissue comprising less than 10% of *C. elegans’* cellular composition (Altun et al. 2009), these observations indicate that ATAC-seq performed on a whole organism is sensitive enough to identify functional tissue-specific enhancers.

We next sought to determine whether ATAC-seq dynamics could be leveraged to identify novel functional enhancers. Previous work has demonstrated that enhancers are precisely activated/inactivated at very specific times in development (Bonn et al. 2012; Kvon 2015; Sun et al. 2015). We hypothesized that the temporal resolution of our ATAC-seq data (due to the precise synchronization of *C. elegans* populations) would enable us to capture distal regulatory dynamics of development. To experimentally test the functional enhancers predicted by our ATAC-seq data, we selected 13 distal (i.e. at least 1kb away from the nearest TSS) non-coding ATAC-seq peaks that exhibited the largest fold-changes in accessibility between any two stages. We generated multiple transgenic *C. elegans* strains with these 13 putative enhancer regions upstream of a minimal promoter (*pes-10*) driving expression of green fluorescent protein (GFP) containing a nucleolar localization signal (NoLS) (Fig 3A, Table 1, Supplementary Tables 12 and 13). To control for specificity, we also tested regions flanking 10 of the 13 putative enhancer regions (Table 1, Supplemental Tables 12 and 13). We created numerous (8-11) independent genetic strains for each validated regulatory site to ensure that no artifact of transgenesis (e.g. hybridization of the *rol-6* marker in the extrachromosomal array) could be driving spurious GFP expression (Table 1). By fluorescence microscopy, we examined the spatiotemporal GFP pattern in these transgenic strains to assess the temporal- and tissue-specificity of the putative enhancers and control regions. Using stringent criteria for defining enhancer activity (see Methods), we found that 6 of the 13 putative enhancer regions (identified by dynamic distal non-coding ATAC-seq peaks) led to specific and consistent spatiotemporal GFP pattern and, for all but one of these, this pattern was observed regardless of the genomic orientation of the region – an important characteristic of enhancers (Table 1, Fig. 3B-G, Supplemental Fig. 5A-E) (Ren and Yue, 2015). In contrast, 0 of the 10 flanking regions led to such a GFP pattern, indicating enrichment for functional enhancers in distal non-coding ATAC-seq peaks that change between stages (p = 0.038, one-sided Fisher’s exact test) (Table 1). Thus, dynamic chromatin accessibility on its own can mark functional enhancers and can be used to successfully identify novel distal regulatory regions.

**Figure 5.**
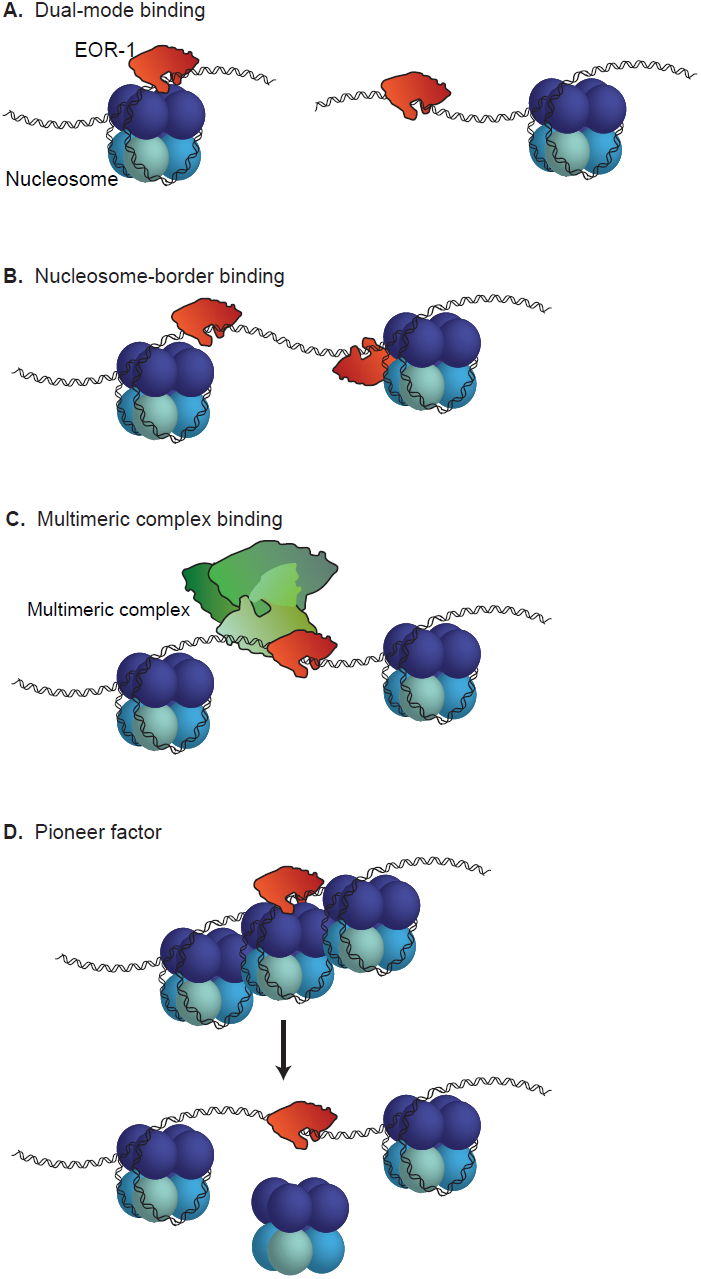
Possible models to explain EOR-1 binding characteristics. EOR-1 could either: (*A*) bind both open and closed chromatin, (*B*) bind immediately adjacent to nucleosomes, (*C*) act as part of a large complex, or (*D*) act as a pioneer factor by binding and contributing to opening closed chromatin.

As expected, the enhancers we experimentally identified display diverse spatiotemporal activity patterns. Three of the enhancer regions (putatively associated with *gei-13*, *mlt-8*, and *nhr-25*) are active in the head or tail hypodermis during development, while others are active specifically in the pharynx (C54G6.3) or along the flank of the worm (*swip-10*). In addition, the enhancers are located at a wide diversity of genomic positions relative to their putatively associated gene: upstream of the TSS (at distances varying from 1-9kb), within introns, as well as downstream of the coding sequence. An interesting example is the *nhr-25-*associated enhancer, which is downstream of the 3’ UTR, more than 5kb downstream of the TSS (Fig. 3D) (this enhancer is also downstream of *npr-10* 3’ UTR on the other side, about 10kb from the TSS). *nhr-25* is a conserved nuclear receptor primarily expressed in the hypodermis (and somatic gonad) during larval development (Gissendanner et al. 2000). The 3’ *nhr-25* enhancer specifically drives GFP expression in approximately 20 hypodermal cells in the head and tail of the worm during larval development (Fig. 3E). The limited expression pattern of the 3’ *nhr-25* enhancer (as well as the other enhancers we identified) indicates that ATAC-seq performed in whole worms is sensitive enough to identify regulatory regions active only in specific cell types, though we cannot rule out the possibility that these regions are also accessible, but their activity repressed (e.g. by H3K27me3) in other cells.

Collectively, these findings demonstrate the power of using ATAC-seq dynamics as an unbiased approach to identify functional and conserved enhancers active in a small subset of cells. When applied to whole organisms or complex samples composed of diverse cellular populations, this approach could capture regulatory regions that may have been missed in studies of isolated cell populations.

### Specific motifs for transcription factors predict changes in chromatin accessibility

To explore the regulatory underpinnings of chromatin accessibility dynamics, including that of enhancers, we examined the occurrence of experimentally defined *C. elegans* transcription factor (TF) binding motifs (Narasimhan et al. 2015) in the dynamic chromatin accessibility regions identified by ATAC-seq. In regions that change chromatin accessibility between early embryo and L3 or between L3 and young adult, we observe significant enrichment of motifs associated with TF homologs and orthologs that have previously been connected to chromatin accessibility dynamics (Fig. 4A; Supplemental Fig. 6A). For example, the motif for BLMP-1, the *C. elegans* ortholog of human BLIMP-1/PRDM1, a TF that recruits chromatin-remodeling complexes in human B cells (Minnich et al. 2016), is enriched in peaks that are more accessible in L3 when compared to embryos or adults. Furthermore, the motif for ELT-3, a GATA TF, is enriched in peaks more accessible in L3 than embryos (the GATA family was recently shown to be an important regulator of chromatin accessibility during human hematopoiesis (Corces et al. 2016)).

The DNA binding motif for EOR-1, which resembles a dimeric version of the canonical GAGA-motif, was significantly enriched in distal non-coding ATAC-seq peaks (Supplemental Fig. 6B) and in ATAC-seq peaks that gained in accessibility in L3 versus embryo (Fig. 4A). This GAGA/EOR-1 motif was also present in 2 of the 5 functional enhancers that we experimentally validated, including the *nhr-25* associated enhancer described above. TFs binding GAGA-motifs have been identified as modulators of chromatin structure dynamics from plants to humans (Hecker et al. 2015; Lund et al. 2013; Srivastava et al. 2013) and the GAGA motif itself is required for full functionality in two previously defined *C. elegans* enhancers (Harfe et al. 1998; Jantsch-Plunger et al. 1994). Finally, GAGA/EOR-1 factors might play an important role in the regulation of chromatin accessibility as EOR-1 genetically interacts with components of two nucleosome remodeling complexes, SWI/SNF and RSC (Lehner et al. 2006), and a dimeric GAGA-motif nearly identical to the EOR-1 motif is highly enriched in ChIP-seq peaks for two separate SWI/SNF components in *C. elegans* (Riedel et al. 2013).

To independently verify the importance of the GAGA/EOR-1 motif, we developed a machine learning model to identify the TF binding motifs that are predictive of regions that gained or lost accessibility between early embryo and larval stage 3. This machine learning method is unbiased and allowed for the integration of the largest available set of previously defined *C. elegans* TF binding motifs (107 high-confidence motifs from 98 TFs out of more than 750 TFs in the *C. elegans* genome (Narasimhan et al. 2015)) as well as 59 motifs discovered *de novo* in regions that changed in accessibility between early embryo and L3 (Fig. 4B; Supplemental Fig. 6C) (see Methods). Using this machine learning framework, we identified the motifs that are the most predictive of ATAC-seq dynamics; the top three most informative motifs in our model are the GAGA-motif that we identified above and motifs for known chromatin regulators (BLMP-1 and ELT-3) (Fig. 4C). Thus, machine learning independently supports the GAGA/EOR-1 motif as a potential important regulator of chromatin accessibility.

We next investigated whether EOR-1 may play a role in chromatin accessibility changes. An EOR-1 ChIP-seq dataset at L3 (Araya et al. 2014) revealed enrichment for a dimeric GAGA-motif (Supplemental Fig. 7A) and was significantly enriched for regions that overlapped ATAC-seq peaks that gain accessibility in L3 compared to early embryo (Supplemental Fig. 7B), indicating that EOR-1 is indeed bound to the GAGA-motif and that EOR-1 may be involved in regulating chromatin accessibility. To examine if EOR-1 is found at closed chromatin, we quantified not only ATAC-seq signal but also ATAC-seq fragment size, as larger ATAC-seq fragments correlate with higher nucleosome occupancy and less accessible chromatin regions (Bao et al. 2015; Buenrostro et al. 2013; Schep et al. 2015). EOR-1 ChIP-seq peak summits at L3 have significantly larger fragment sizes (indicative of less accessible chromatin regions) than all 24 canonical TFs and the histone deacetylase HDA-1 ChIP-seq peak summits (FDR < 0.05) (Fig. 4D; Supplemental Fig. 7C, 7D; Supplemental Table 14). In fact, the only factor with larger ATAC-seq insert sizes was the heterochromatin-associated protein HPL-2. The larger ATAC-seq insert sizes at EOR-1 binding sites are unlikely to be due to the binding of other TFs hindering the ability of transposons to interact with the genomic DNA because EOR-1 ChIP-seq peaks did not show unusual overlap with the other TFs assessed (Supplemental Fig. 7E). These observations are consistent with the possibility that EOR-1 is uniquely present at regions with less accessible chromatin compared to other TFs.

To more closely investigate the chromatin accessibility landscape of EOR-1 peaks at the L3 stage, we generated aggregated histograms of the median ATAC-seq fragment sizes for the ChIP-seq peak summits of EOR-1, all canonical TFs, and the chromatin-associated factors HPL-1 and HDA-1 (Fig. 4E; Supplemental Fig. 7F). As expected, canonical TFs are predominantly found at regions with short ATAC-seq insert sizes (less than 147 bp, the length of DNA wrapped around a nucleosome), whereas chromatin factors, especially HPL-2, are also present at regions with larger ATAC-seq fragment sizes (with many greater than 147 bp). Strikingly, EOR-1 was found both at regions with short ATAC-seq insert sizes as well as at regions with larger ATAC-seq fragment sizes (greater than 147 bp). Together, these results suggest that EOR-1 may in some cases, bind to or at least immediately adjacent to nucleosome-occupied DNA.

We next quantified nucleosome occupancy dynamics in the summits of ChIP-seq peaks using publicly available histone H3 ChIP-seq data (Contrino et al. 2012; Gerstein et al. 2010). EOR-1 peak summits exhibited a large decrease in nucleosome occupancy from embryo to L3, and this was distinct from the canonical TFs assessed (Fig. 4F). In contrast, chromatin factors such as HPL-2 and HDA-1 exhibited an increase in nucleosome occupancy. These results suggest that EOR-1 binding sites exhibit changes in chromatin accessibility during the transition from early embryos to L3 that are unique from the other factors analyzed.

Taking these observations collectively, we propose four potential models to explain the unique binding profile of EOR-1: i) EOR-1 is capable of binding to both open and closed chromatin (depending on the loci or the tissues) (Fig. 5A); ii) EOR-1 binds immediately adjacent to nucleosomes (resulting in larger ATAC-seq fragment sizes), including those nucleosomes that shift location between early embryo and L3 (explaining the change in accessibility and nucleosome signal) (Fig. 5B); iii) EOR-1 binds open chromatin as part of a larger complex including factors not assayed here (e.g. EOR-2 (Howell et al. 2010)), thereby hindering the ability of transposons to interact with the genomic DNA (Fig. 5C); and iv) EOR-1 binds closed chromatin as a pioneer factor and contributes to its opening (Fig. 5D). Future experiments, such as nucleosome binding assays and chromatin profiling of *eor-1* mutants, will be needed to distinguish between these models and to fully elucidate the mechanism underlying the unusual binding profile of EOR-1. Collectively, these analyses suggest that ATAC-seq on a complex sample can identify factors with unusual chromatin binding patterns that could regulate chromatin dynamics during development.

## Discussion

Here we show for the first time that sensitively measuring chromatin accessibility in a whole organism can detect dynamic changes throughout development and even identify novel functional enhancers active in only a small subset of the whole organism. Between the 3 developmental stages surveyed, we identified over 30,000 accessible sites which could serve as a catalogue to facilitate the discovery of previously unknown distal regulatory loci (Supplemental Table 3) such as insulators and enhancers. This approach, developed here for *C. elegans*, should be readily applicable to other complex samples *in vivo* such as whole mammalian organs or tumor samples.

In *C. elegans*, enhancer identification has historically been limited, with most previous studies employing a single gene approach and focusing on promoter-proximal regions (Harfe et al. 1998; Jantsch-Plunger et al. 1994; Lei et al. 2009). Others have attempted to identify enhancers genome-wide, but have not functionally validated predicted enhancer activity (Chen et al. 2013; Vavouri et al. 2007). In this study, we identify and functionally characterize distal-ATAC-seq peaks as novel, active enhancers. These enhancers have a range of spatiotemporal activity patterns that are orientation-independent, and are found at a diversity of genomic locations, suggesting that enhancers may be more prevalent in *C. elegans* gene regulation than previously appreciated. An interesting example is the enhancer downstream of the conserved transcription factor *nhr-25*. This putative-*nhr-25* enhancer is located 5kb downstream of the *nhr-25* TSS and acts in an orientation-independent manner. Thus, this enhancer might be looping to the *nhr-25* TSS to enhance transcription, although the limited spatial resolution of current Hi-C data in *C. elegans* (Crane et al. 2015) does not provide information on the three-dimensional chromatin structure of the *nhr-25* locus. This enhancer model contrasts with the promoter-proximal model most often thought to regulate gene expression in *C. elegans*, but agrees with the recent Hi-C study which identified insulator-like loci throughout the *C. elegans* genome (Crane et al. 2015). Insulators are important regulators of three-dimensional chromatin architecture (Schoborg et al. 2014). Interestingly, *nhr-25* orthologs in fly (Ftz-F1), mouse (NR5A), and human (NR5A), exhibit a consistent signature of 3-dimensional chromatin architecture downstream of the 3’ UTR (insulator class I in flies, and CTCF-binding sites in mice and humans (Attrill et al. 2016; Rosenbloom et al. 2013; Sharov et al. 2006)). While *C. elegans* does not possess a CTCF ortholog, our results raise the intriguing possibility that the *nhr-25* enhancer we have identified is evolutionarily conserved.

Furthermore, we have uncovered a potential role for a likely GAGA factor in *C. elegans*, EOR-1. The EOR-1 motif we initially detected closely resembles a dimeric version of the canonical GAGA-motif bound by Trl/GAGA-Associated Factor (GAF) in *Drosophila*. GAF is a multi-faceted transcription factor that can associate with heterochromatin (Raff et al. 1994), remodel chromatin in concert with nucleosome remodelers (Okada et al. 1998), and acts as a transcriptional activator in part due to its ability to increase chromatin accessibility (Adkins et al. 2006). Like *Drosophila* GAF, *C. elegans* EOR-1 is a transcriptional activator in the Ras/ERK signaling pathway that has been suggested to also repress gene expression (Liu et al. 2011a). This dual action (gene activation and repression) is particularly interesting considering the bimodal binding pattern we found when examining ATAC-seq fragment size in EOR-1 binding sites. GAF and EOR-1 are also similar in that both proteins have a BTB/POZ domain on their N-terminal as well as C2H2 zinc-fingers and polyQ domains on their C-terminals (Howard et al. 2002).

In *C. elegans*, EOR-1 genetically interacts with at least two chromatin remodeling complexes (SWI/SNF and RSC) (Lehner et al. 2006). This is noteworthy given our findings that EOR-1 is found in less accessible and potentially even nucleosome-occupied regions of the genome, and that the EOR-1 motif is predictive of increased accessibility in development. GAGA-factors are conserved regulators of gene expression (Mahmoudi et al. 2002; Ohtsuki et al. 1998; Petrascheck et al. 2005; Srivastava et al. 2013), and a GAGA motif has been found to be necessary for enhancer functionality in *C. elegans* (Harfe et al. 1998). Indeed, 2 out of 5 of the novel validated enhancers identified in this study (*nhr-25*, and *swip-*10) contain an EOR-1/GAGA motif. Collectively, these data suggest that EOR-1 might regulate chromatin accessibility at enhancers in *C. elegans* and potentially other species.

Using *C. elegans* as a paradigm, we have shown that ATAC-seq performed on complex, heterogeneous samples can reveal novel, spatiotemporally-specific genetic regulators, and that measuring chromatin accessibility across a developmental time course can identify important dynamic regions. We have highlighted important applications of this approach: discovering functional distal regulatory regions active in only a small subset of the total sample and identifying candidate regulators of genome-wide chromatin dynamics. This dataset, which represents an initial atlas of genome-wide chromatin accessibility and candidate distal regulatory sites in *C. elegans*, should be a valuable resource for the community. The fact that a genome-wide chromatin accessibility assay performed in whole organisms can sensitively identify previously undiscovered functional enhancers *in vivo* raises the exciting possibility that distal regulation plays a more important role than previously believed in the nematode.

## Methods

Brief methods can be found below, but in all cases more details can be found in the supplemental methods.

### *C. elegans* ATAC-seq

Three sets of completely independent biological replicates were prepared by harvesting and then flash-freezing tightly temporally-synchronized samples at three different stages: early embryo (*in utero*), larval stage 3 (36 hr post egg lay), and young adult (57 hr post egg lay). Native nuclei were purified from frozen samples using mechanic homogenization as previously described (Haenni et al. 2012). The purified nuclei were immediately used for the ATAC-seq protocol (Buenrostro et al. 2013). An input control was also generated by using 10 ng of genomic DNA. Sequencing was performed using 101 bp paired-end sequencing on an Illumina Hi-seq 2000.

### ATAC-seq alignment and peak calling

The ATAC-seq libraries were sequenced to a median depth of over 17 million unique, high-quality mapping reads per sample (Supplemental Table 1). Prior to mapping, standard next generation sequencing quality control steps, as well as ATAC-seq specific quality control steps were performed. Individual replicate ATAC-seq peaks were called using a custom pipeline which used MACS (Zhang et al. 2008) (v2.1) to call the peaks. To identify the most high-confidence set of peaks, we employed a strategy that emphasized peaks which were consistently observed across replicates. All peaks were masked for regions with significant signal in the input control and for previously identified regions known to give spurious results in next generation sequencing assays (Boyle et al. 2014). The result was a set of 30,832 consensus ATAC-seq peaks.

### Selection of putative enhancers and generation of enhancer reporter constructs

For the enhancer screen, each putative regulatory region was cloned in the pL4051 plasmid (a gift from Andrew Fire), upstream of a minimal promoter (*pes-10*) driving expression of a *C. elegans* intron- and photo-stability-optimized GFP containing an N-terminal nucleolar localization signal (NoLS). Putative regulatory regions were chosen by selecting ATAC-seq peaks that exhibited the largest differential accessibility between two stages and that were at least 1kb from a transcription start site. Flanking negative control regions were chosen by selecting regions within 2kb of the putative regulatory regions that were not in peaks of accessibility. Primers were designed to amplify each region as well as 50-500bp flanking either side (Supplemental Table 13).

### Enhancer screen in *C. elegans*

Stable extrachromosomal transgenic lines for putative enhancer regions (in both orientations), negative control regions, and the no-insert control were generated. Multiple independent lines were generated per construct (Table 1). For each line, mixed-staged worms were screened for GFP signal distinct from the no-insert control background signal, which is 1-2 nuclei near the pharynx in all larval and adult stages (Supplemental Fig. 5E). To quantify the consistency of GFP expression pattern, all lines (including the negative control lines) (Table 1) were scored for GFP expression pattern in a blinded manner.

### Machine learning models to predict accessibility changes with motifs

To predict changes in accessibility between early embryo and larval stage 3, the number of each mapped *C. elegans* motif from cisBP (v1.02) (Ray et al. 2013) and each *de novo* discovered motif found in the dynamics ATAC-seq peaks between early embryo and L3 (59 in total) was counted to create a matrix of 166 motif-counts and 30,832 ATAC-seq peaks. The ATAC-seq peaks were split into a training set (70%) and a testing set, and several classification models evaluated. We found that a generalized boosting model (GBM) (Ridgeway 2015) had the best accuracy while still allowing for interpretation of which motifs were the most informative. Given the unbalanced classification problem, we used balanced accuracy as our primary metric of classification success (Supplemental Fig. 6D).

## Data Access

Raw reads as well as stage-specific peaks can be found on the Gene Expression Omnibus website using accession GSE89608. All code can be found at https://github.com/brunetlab/CelegansATACseq.

## Acknowledgements

We thank Dr. Elizabeth Noblin and Max Lenail for their help imaging enhancer reporter strains, and Dr. Eric Greer for assistance in developing the nuclei isolation protocol. We thank Dr. Elizabeth Noblin, Dr. Lauren Booth, and Dr. Bé ré nice Benayoun for critically reading the manuscript. We thank Dr. Bé ré nice Benayoun for feedback on computational pipeline and analysis, Daniel Kim for use of his ‘smartMerge’ script, and Dr. Andy Fire and Dr. Joanna Wysocka for suggestions on the project. Supported by NIH DP1 AG044848 (A.B.), a seed grant from the Discovery Innovation Fund (A.B.), and NSF graduate fellowship (A.C.D.).

## Author contributions

A.C.D. and A.B. planned the study. A.C.D. performed the ATAC-seq experiments, designed the analytical pipeline, analyzed and interpreted the data, and wrote the manuscript with help from A.B. R.Y. generated the transgenic enhancer reporter lines, imaged and analyzed GFP expression, and led revisions to the manuscript. A.K. provided intellectual guidance with ATAC-seq analysis and machine learning. J.D.B. and W.J.G. provided early access to ATAC-seq protocols and feedback on the design. All authors discussed the results and commented on the manuscript.

---

**Supplemental Figure 1.**
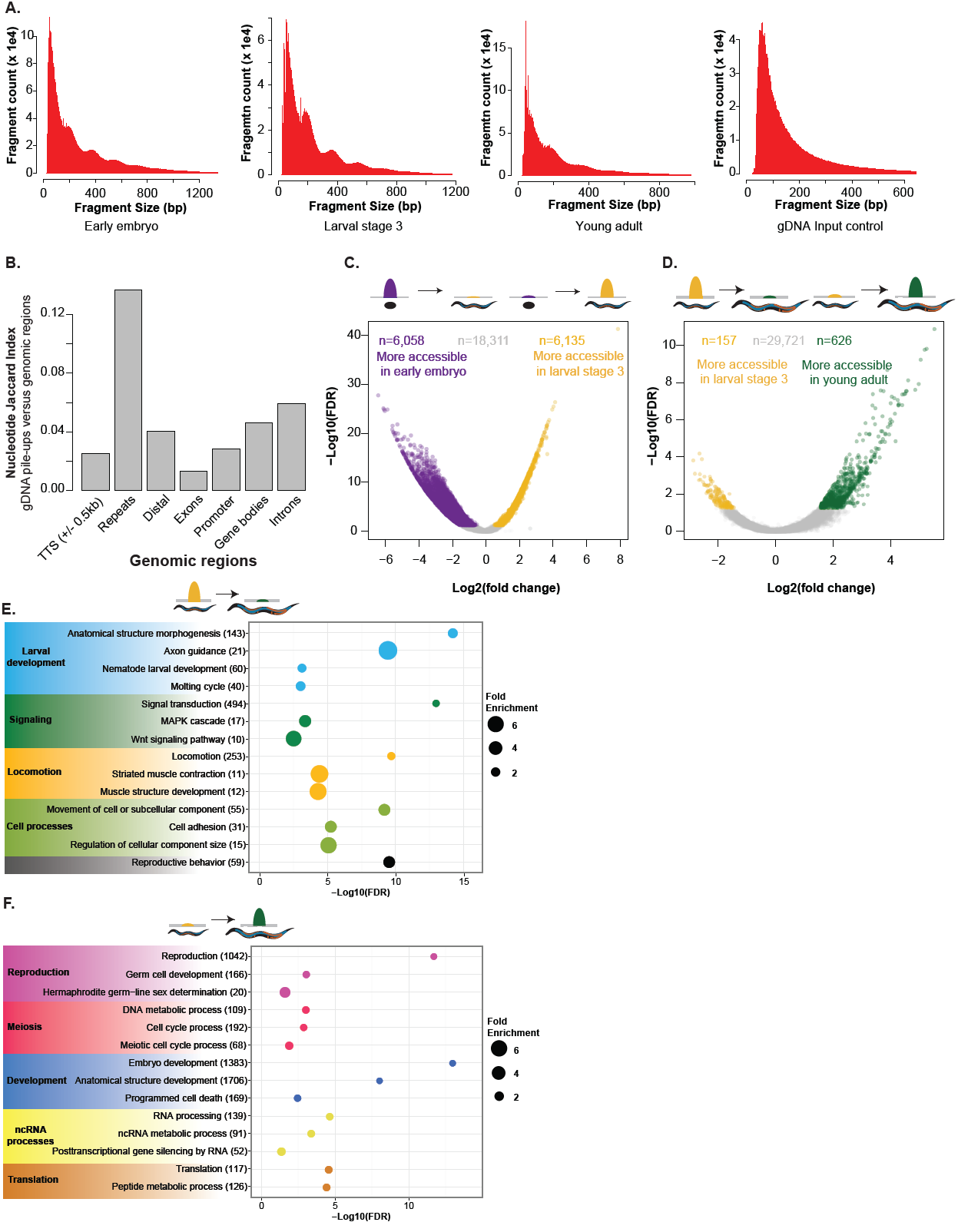
(*A*) Histograms of fragment size for representative experimental replicates (first three panels from the left) show consistent approximately 147bp periodicity, likely corresponding to nucleosome-protected fragments. This periodicity is not present in the purified genomic DNA (gDNA) input control (far right panel). (*B*) Read pile-ups in genomic DNA input control overlap annotated repeats. The single base pair Jaccard index (a measure of nucleotide overlap) was calculated for the called gDNA read pile-ups versus genomic locations (downloaded from the UCSC Genome Browser). (*C,D*) Significantly differential consensus ATAC-seq peaks between early embryo (embryo) and larval stage 3 (L3) (C), and larval stage 3 (L3) and young adult (adult) (*D*); FDR < 0.05). (*E,F*) Genes that lose accessibility between L3 and adult are enriched for larval development functions (E), while genes that gain accessibility are enriched for adult-related functions (F); all calculations and genes lists are from GOrilla and the number of genes contributing to the enrichment of each term are listed in parentheses

**Supplemental Figure 2.**
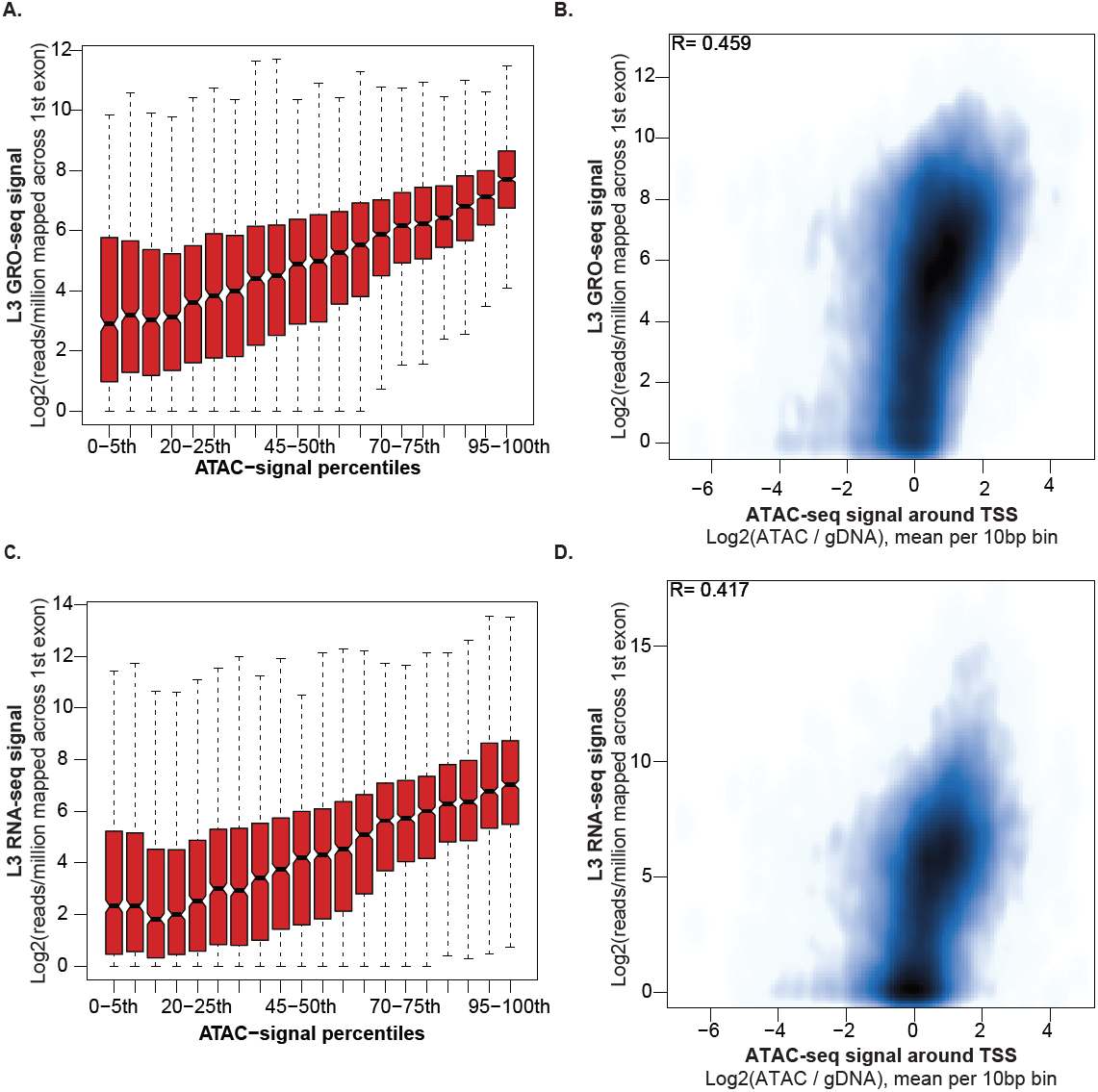
(*A*) Interquartile bar plots illustrating transcriptional activity with increasing ATAC-seq signal (binned as 5 percentile ranges). For each gene, the ATAC-seq signal (+/- 1kb around the TSS) and nascent transcription (RPKM across RefSeq first exons) was calculated using L3 ATAC-seq data and publicly available L3 GRO-seq data. (*B*) Smoothed scatterplot illustrating the same relationship between TSS accessibility and nascent transcription (Pearson r is shown). (*C, D*) Interquartile bar plot and smoothed scatterplot illustrating the relationship between TSS accessibility and transcriptional activity using publicly available RNA-seq data at the L3 stage

**Supplemental Figure 3.**
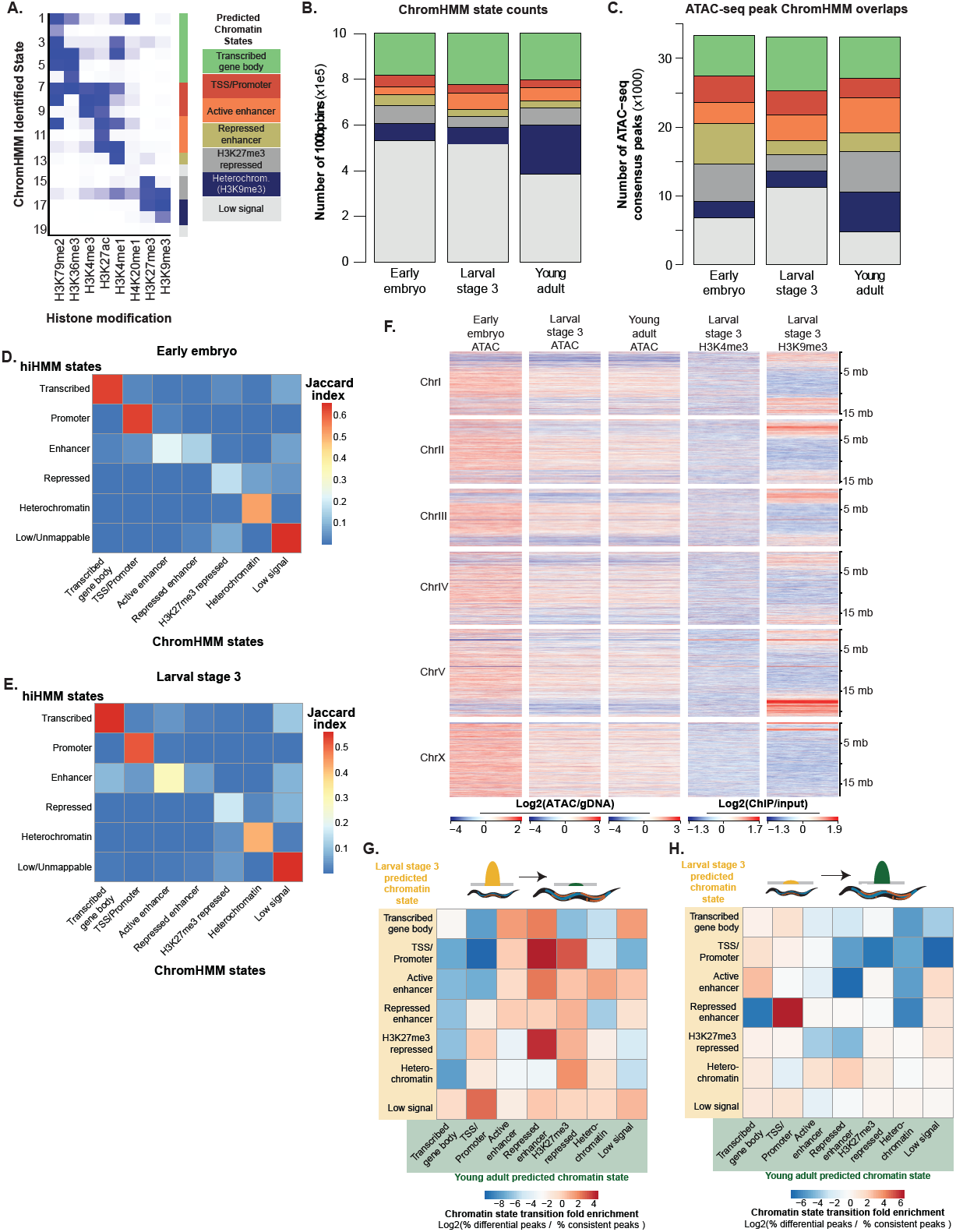
(*A*) Heatmap of emission parameters from ChromHMM show enrichment of the histone modifications in each ChromHMM-predicted chromatin states as well as the 7 core chromatin states. Each of the 19 original states was assigned to a core chromatin state using canonical associations of histone modifications. (*B*) The number of 100bp bins (the resolution used for building the ChromHMM model) found in each predicted state for each of the three stages. (*C*) The number of ATAC-seq peaks which overlapped by at least half their length in each stage-specific ChromHMM predicted state. (*D,E*) ChromHMM predicted chromatin states closely resemble predictions made by (Ho et al. 2014) using hiHMM a method similar to ChromHMM. For both early embryo (D) and larval stage 3 (E) the Jaccard index (a measure of nucleotide overlap) for each of the 7 ChromHMM-predicted states was calculated for each of the 6 chromatin states predicted by Ho et al. 2014 using hiHMM. (*F*) ATAC-seq signal is highest in the gene-dense center of *C. elegans* chromosomes and notably anti-correlated with heterochromatin-associated H3K9me3 ChIP-seq. (*G,H*) Decreases in accessibility in ATAC-seq peaks are enriched for transitions from active regulatory to inactive chromatin states (G), while increases in accessibility are enriched for transitions from inactive chromatin states to active regulatory states (H).

**Supplemental Figure 4.**
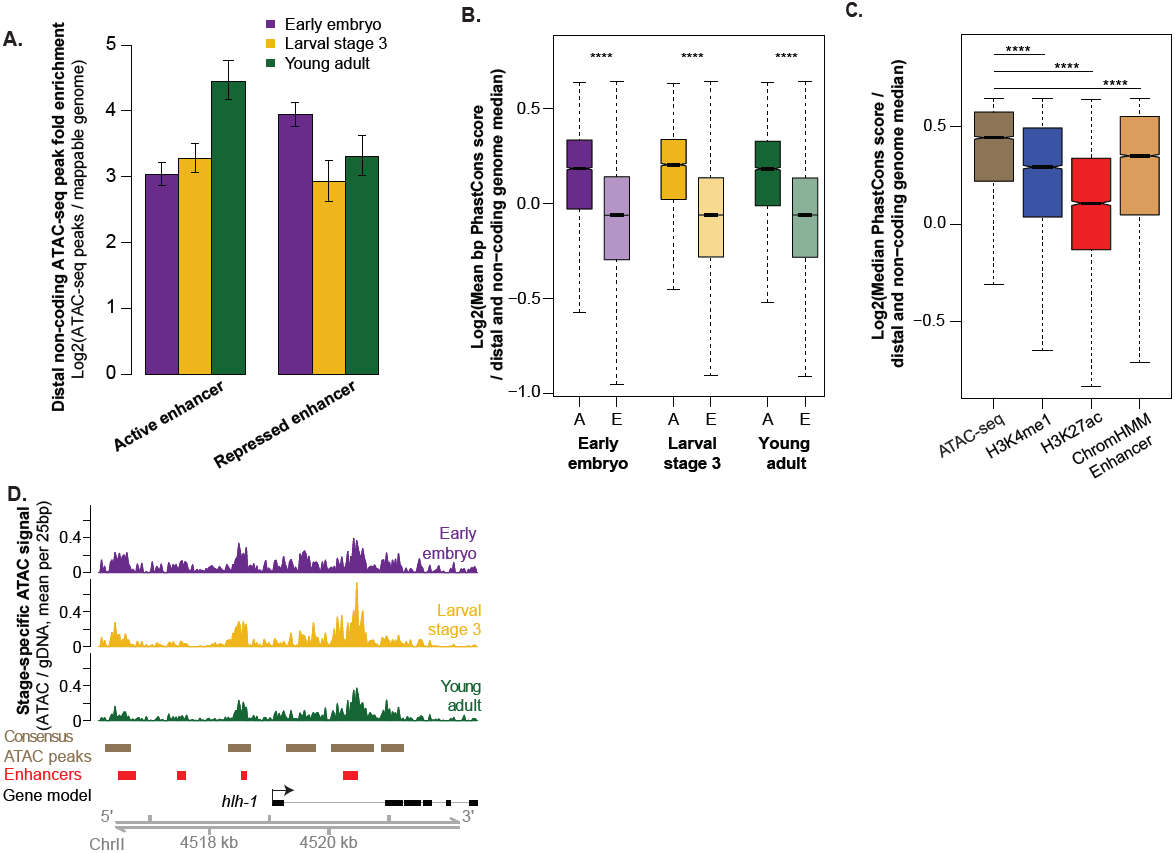
(*A*) Distal non-coding ATAC-seq peaks are enriched for ChromHMM-predicted enhancer chromatin states. Significance was assessed using 10,000 bootstrap iterations and error bars represent 95% confidence intervals (all enrichments p < 1e-4). (*B*) The median single base pair conservation score for every stage-specific distal non-coding ATAC-seq peak was normalized to the genome-wide distal non-coding median (A: Actual, darker hues) and compared to expected scores (E: Expected, light hues) derived from randomizing peak locations across the distal non-coding genome (^***^ p < 1e-323, Kolmogorov-Smirnov (KS) test). (*C*) Consensus distal non-coding ATAC-seq peaks are more highly conserved than distal non-coding histone modification peaks often associated with enhancers. Pooled H3K4me1 and H3K27ac ChIP-seq peaks are from all three stages. ChromHMM enhancers are regions predicted to be in an active or poised enhancer state in any of the three stages assayed. ^***^ p < 1e-323, KS test. (*D*) Consensus ATAC-seq peaks overlap experimentally defined enhancers near the MyoD homolog, *hlh-1*. Functionally defined enhancers (red) were from (Lei et al. 2009).

**Supplemental Figure 5.**
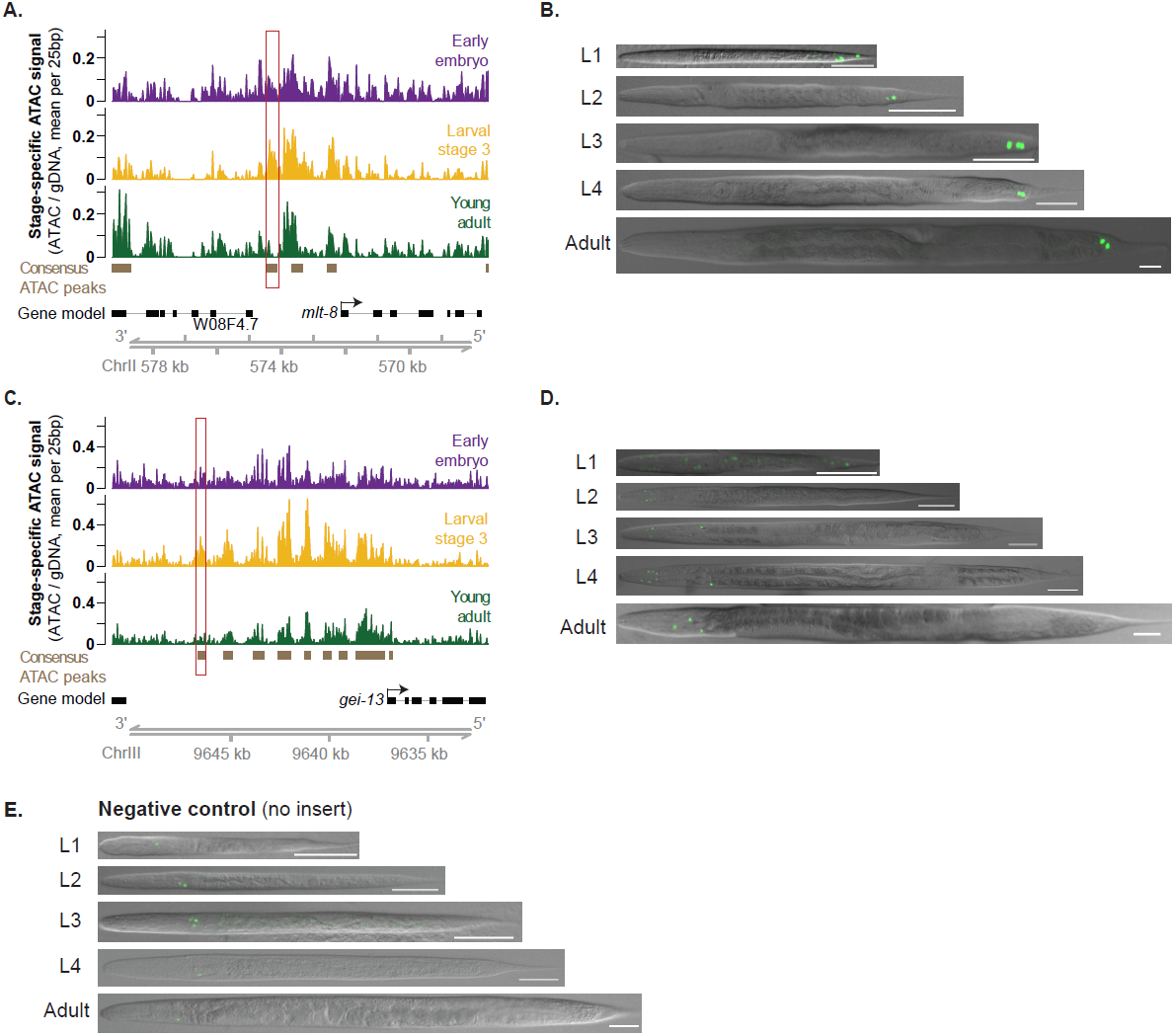
(*A,C*) ATAC-seq signal near *mlt-*8 and *gei-13* including the regions tested for enhancer activity (boxed in red). Note both plots are in reverse orientation for consistency with Figure 4. (*B,D,E*) Representative images of staged *C. elegans* transgenic lines for *mlt-8* (B), *gei-13* (D) and no-insert negative controls (E). The scale bar is 50 μm, and all images were straightened with ImageJ and are greyscale images with florescence overlaid.

**Supplemental Figure 6.**
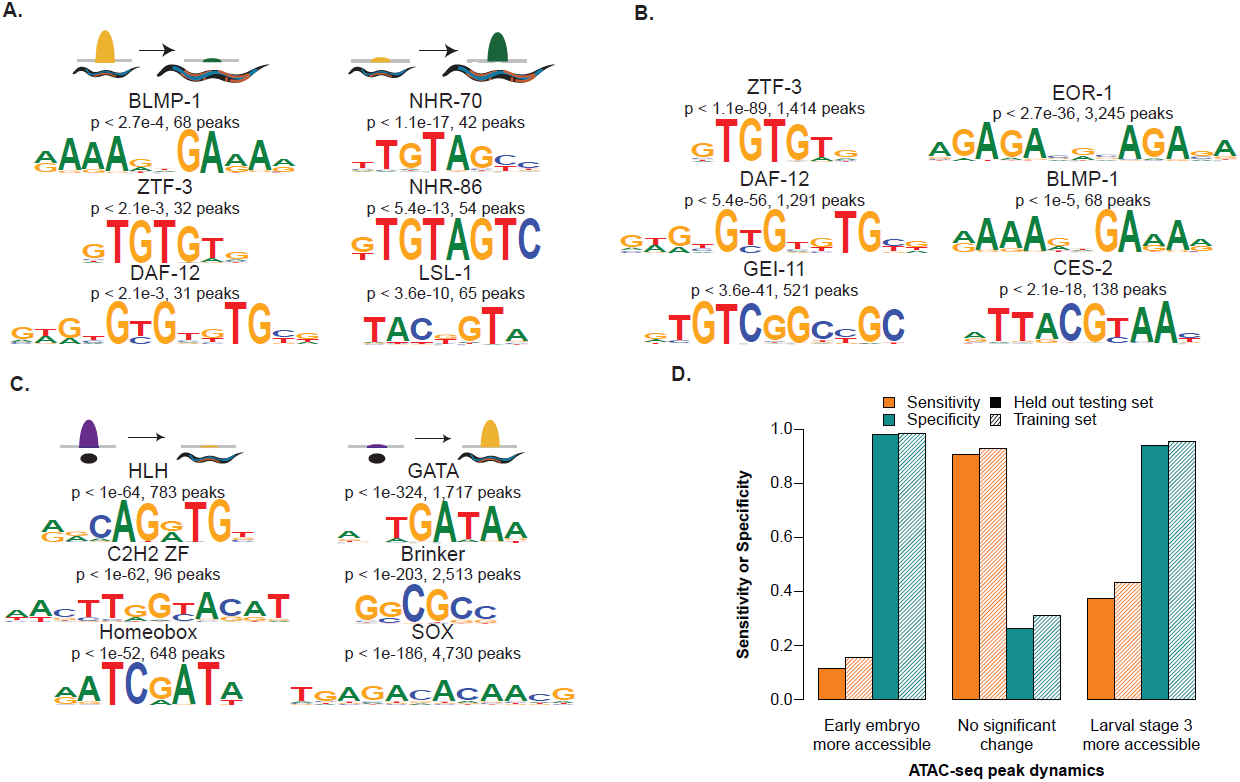
(*A*) ATAC-seq peaks which decreased (left) or increased (right) accessibility between L3 and young adult are enriched for previously identified transcription factor binding motifs; p-values are Benjamini-Hochberg corrected for multiple hypothesis testing. (*B*) Enrichment of experimentally defined *C. elegans* transcription factor motifs in distal non-coding ATAC-seq peaks; p-values are Benjamini-Hochberg corrected for multiple hypothesis testing. (*C*) *de novo* motifs identified in peaks which were decreased (left) or increased (right) between embryo and L3. Motifs were discovered using Homer and a background set of all unchanging ATAC-seq peaks. (*D*) Evaluations of predictions of chromatin accessibility changes between L3 and early embryo. The sensitivity (True Positive Rate) and specificity (True Negative Rate) for both testing and training sets of peaks were determined for each dynamic ATAC-seq peak type.

**Supplemental Figure 7.**
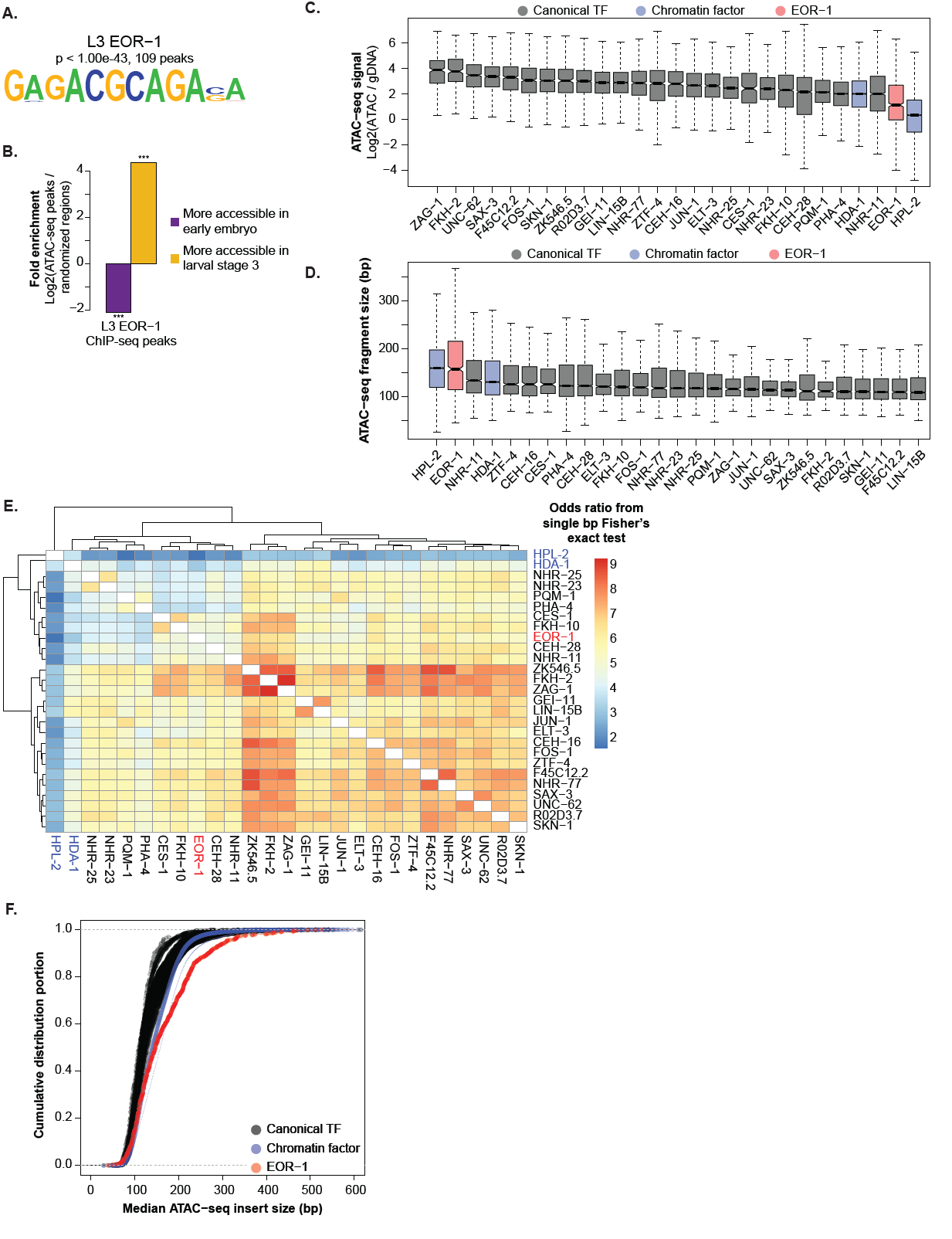
(*A*) The most significantly enriched *de novo* motif in the L3 EOR-1 ChIP-seq peaks. (*B*) L3 EOR-1 ChIP-seq peaks are significantly enriched for ATAC-seq peaks that increase in accessibility between early embryo and L3. Fold enrichment and significance was calculated by comparison to a null distribution of 10,000 iterations. (*C,D*) L3 EOR-1 ChIP-seq peak summits are less accessible than other L3 canonical TF and chromatin factor ChIP-seq peak summits. For each TF ChIP-seq, ATAC-seq signal (C) and fragment size (D) within the midpoint (+/- 50 bp) of every peak is reported. (*E*) Heatmap illustrating the co-localization of L3 ChIP-seq peaks overlapping the peaks of other factors. Odds Ratio calculated from Bedtools’ implementation of Fisher’ s exact test, which uses the entire genome as background. *(F)* The cumulative distribution functions for the median L3 ATAC-seq fragment size at the midpoint (+/- 50 bp) of L3 ChIP-seq peak summits represented in Figure 4E.

